# Mutation signatures reveal biological processes in human cancer

**DOI:** 10.1101/036541

**Authors:** Kyle R. Covington, Eve Shinbrot, David A. Wheeler

## Abstract

Replication errors in the genome accumulate from a variety of mutational processes, which leave a history of mutations on the affected genome. The relative contribution of each mutational process has been characterized by non-negative matrix factorization and has lead to deeper insight into both mutational and repair processes contributing to cancer. However current implementations of NMF have left unresolved some specific patterns that should be present in the mutation data and have not generated signatures designed for classification. Here, we use a variant of NMF, termed non-smooth NMF, to generate sparse matrix factorizations of somatic mutation profiles present in 7129 tumors. nsNMF factorization revealed 21 mutational signatures. We found three APOBEC mutational processes clearly segregating with the published APOBEC enzymology and trans-lesion repair processes. We discovered several signatures differed between geographic locations even between closely related tissues.

## Introduction

Understanding the mutational mechanisms in cancer can yield further insights both into the biology of cancer cells, and of cellular processes in general regulating DNA damage and repair. As the vast majority of mutations arising in any given tumor are neutral in function (so-called passenger mutations), the study of mutations arising in tumor data is largely unbiased by phenotypic selection. It is therefore possible to use statistical learning techniques to extract unbiased mutation signatures from raw somatic mutation data. As the data are strictly positive, both in terms of the observed mutations and the effect of any particular mutagen, an approach termed non-negative matrix factorization (NMF) has become a popular method of signature identification [1, 2, 3, 4, 5, 6]. Several studies have reported mutational spectra in cancer cohorts and contrasted differences between cancer types [7, 5, 6] using NMF to decompose the mutation spectrum within each cancer sample. This work has been quite fruitful, identifying both endogenous, such as APOBEC [5, 8, 9] and POLE [10], and exogenous, such as UVB [5], mutation processes.

NMF decomposition is attractive for many reasons including being sensitive to subgroups within the data and allowing classification of new samples. However, different NMF strategies should be employed for different purposes. When attempting to account for the majority of mutations present within the data, it is desirable to minimize the residual error of the solution. On the other hand, using NMF to build classifiers requires minimizing correlations between signatures (increased orthogonality) and reducing noise (increasing sparseness). Most existing mutation signature NMF solutions have been tuned to reduce residual error, at the expense of sparseness and orthogonality [11, 5, 12, 13].

Our objective was to generate mutational signatures for use as a classification tool, such that new datasets could be classified and investigated. We chose a variant of NMF which uses an internal smoothing matrix to drive sparseness (suppress noise) termed non-smooth NMF (nsNMF) [4].

## Results

### Comparison of NMF and nsNMF signatures

We generated a 21 signature solution based on our selection and optimization criteria (see methods) using nsNMF [4], further referred to as the nsNMF signature. We compared the properties of the nsNMF solutions to those previously reported by Alexandrov *et al.* [5] using modifications of the NMF algorithm described by Brunet *et al.* [2], further referred to as the Brunet signature. We examined the orthogonality of the nsNMF and Brunet signatures using cosine similarity (Figure 1A). The average cosine similarity for the Brunet's signature was 0.34 compared to 0.07 for the nsNMF signature, with only 3.8% (16 / 420) of the nsNMF pairings having a cosine similarity greater than 0.34. This “overlapping!” effect is common in NMF and was one of the major motivations for the development of nsNMF [4].

The high correlations in signature solutions within the Brunet signatures effects their utility as classifiers. We generated signature solutions for a set of 485 liver cancer samples published as part of a collaborative effort between the International Cancer Genome Consortia (ICGC) and The Cancer Genome Atlas (TCGA) by Totoki *et al.* [14]. Liver cancers represent an excellent test case for mutation signatures. In addition to being one of the most common cancer types, several mutational processes operate in liver cancer identified by both the Brunet and nsNMF solutions and these cancers have nearly an average mutation rate compared to other cancers. To test the stability of the signature solutions in liver cancers, we distorted the signatures by adding to each case a single mutation (the effect of calling an extra variant) and generated signature solutions from the distorted data. The original and distorted signature solutions were then compared for both cosine similarity and absolute difference (see supplemental information). nsNMF was more resilient to distortion compared to Brunet signatures (Figure 1B, Supplemental Information). Overall, signature distortion was lower for nsNMF with higher cosine similarity values in 55% and a lower absolute distortion in 70% of all mutation distortions (Supplemental information). These effects were much more pronounced in the “dominant” signatures (those accounting for > 15% of the mutation burden in > 10% of the samples) with 65% of distortions favoring nsNMF on the basis of cosine similarity and 82% favoring nsNMF on the basis of absolute distortion. We should point out that as the nsNMF solution contains 21 signatures compared to 27 for the Brunet solution, the cosine similarity results are biased in favor of the Brunet solution. In summary, these data indicate that the nsNMF signatures generate more stable signature solutions in cancer samples compared to Brunet signatures.

As expected, mutation signature solutions derived from nsNMF contained many signatures previously associated with identified mutational processes (Figure 2A). For example, signature 6 is highly enriched for C>T mutations at 5'NCG sites and represents the most common signature across all cancer types (Figure 2B). We found signature 14 to be very similar to the mutation pattern induced by UVB radiation and this signature was by far the most penetrant signature in melanoma cancers (Figure 2A and B). Signature 19 was very similar to the POLE-exonuclease mutant samples of the “class A”, active, type [10]. We note that the POLE signature is highly penetrant and some form of POLE existed in nearly every spectral optimization we performed.

Alexandrov *et al.* [5] and Nik-Zainal *et al.* [15] each proposed two likely APOBEC signatures, both of those signatures showed high C>T and C>G in the 5'TCW context with only minor differences in scale between them. We also found two nsNMF signatures consistent with APOBEC mutation (signatures 4 and 9), however, nsNMF has clearly separated the C>T and C>G effects with a cosine similarity of **3.4e-64,** indicating that these signatures are likely the result of separate mutational processes. In fact, two distinct abasic site repair mechanisms exist for the repair of APOBEC-induced cytosine deamination [8]. APOBEC enzymology would predict a third APOBEC mutation signature, as APOBEC3G has a preference for deamination of cytosine 3' of a cytosine (5'CC) as opposed to 3' of thymine [9]. nsNMF was able to identify such a CCW context signature (signature 21) consistent with the activity of APOBEC3G. These findings highlight the ability of nsNMF to isolate minor yet consistent components from complex mixtures.

### Mutation signatures in cancer

Several mutation patterns were observed to be enriched in specific organ environments (Figure 2C). For example, signatures 2, 5, and 16 were all enriched in lung cancer samples. Signatures 2, 5, and 16 all contain a high proportion of C>A mutations consistent with oxidative damage associated with tobacco smoke. Signature 7 was associated with stomach and esophageal cancers and may be caused by exposure to arsenic [16]. We found signature 13 to be highly specific for liver cancers. These data support the concept that many mutational processes are environmental in origin.

### Mutation signatures differ within a cancer

We have applied our signatures to several novel datasets to study the contribution of different mutational mechanisms in cancer biology.

We generated mutation signature solutions for a set of 485 liver cancer samples with sufficient mutation counts to perform signature assignment. These samples and their mutation data had been previously published [14]. Signatures 5, 6,13, and 18 showed a high contribution to mutations in this set (Figure 3A). As shown above, signature 6 is very common across many cancers and represents CpG mutations. Signature 13 was highest in liver cancers in the training data and it's role as a liver selective signature is confirmed by these data. Signature 18 was correlated with high mutation rate (Pearson's correlation; p < 0.0003, Supplemental information) and may represent mutations caused by aristolochic acid exposure [17, 18], which can generate mutations in the liver [19]. Signature 5 showed a strong correlation with race, with only Caucasian and Asians in the United States showing no difference between the races (Figure 3B). Signature 5 was also associated with differences in outcomes (Cox proportional hazards; p < 0.01, Figure 3C) in univariate models, and showed a trend with outcomes in models including race as a covariate (Cox proportional hazards; p < 0.1). The reduction in significance is likely because Asian race was significantly associated with good outcomes in this study (Cox proportional hazards; p < 0.02).

We then used the nsNMF signatures to explore differences in mutation patterns associated with polymerase **-E** (POLE) mutations in colorectal cancers. POLE mutation has been associated with ultra-mutation rate cancers arising in many organs, but predominantly endometrial and colorectal cancers [10]. Shinbrot *et al.* [10] demonstrated a context specific mutation pattern of POLE-exonuclease domain mutant tumors which is identical to Signature 19. We generated mutation signature estimates for 449 colorectal samples within the TCGA colorectal study (Figure 4A). The most prevalent mutation signature was signature 6, which, as described above, is the CpG C>T mutation signature. While signature 6 is the most common mutation type in cancer, colorectal cancers (along with esophageal, stomach, and uterine cancers) were one of the major contributors of signature 6. Signature 19 was highly penetrant in a subset of cancer samples, all of which contained mutations in POLE-exonuclease domain. Signature 19 accounted for at least 40% of all mutations in POLE-exonuclease domain mutant samples (Figure 4B). POLE-exonuclease domain samples showed the highest mutation counts, though, POLE-exonuclease domain mutant samples were still the highest mutation count samples even after subtracting signature 19 mutations away from the total mutations. This would indicate that other mutational processes could also be present in these samples. Mutation signatures also differed between colon and rectal samples with respect to signature 6 (Figure 4C) and all signatures were different between MSI and MSS samples (Figure 4D).These findings indicate that the nsNMF mutational signatures are associated with both environmental and genetic mutagen exposure.

**Figure 1:**
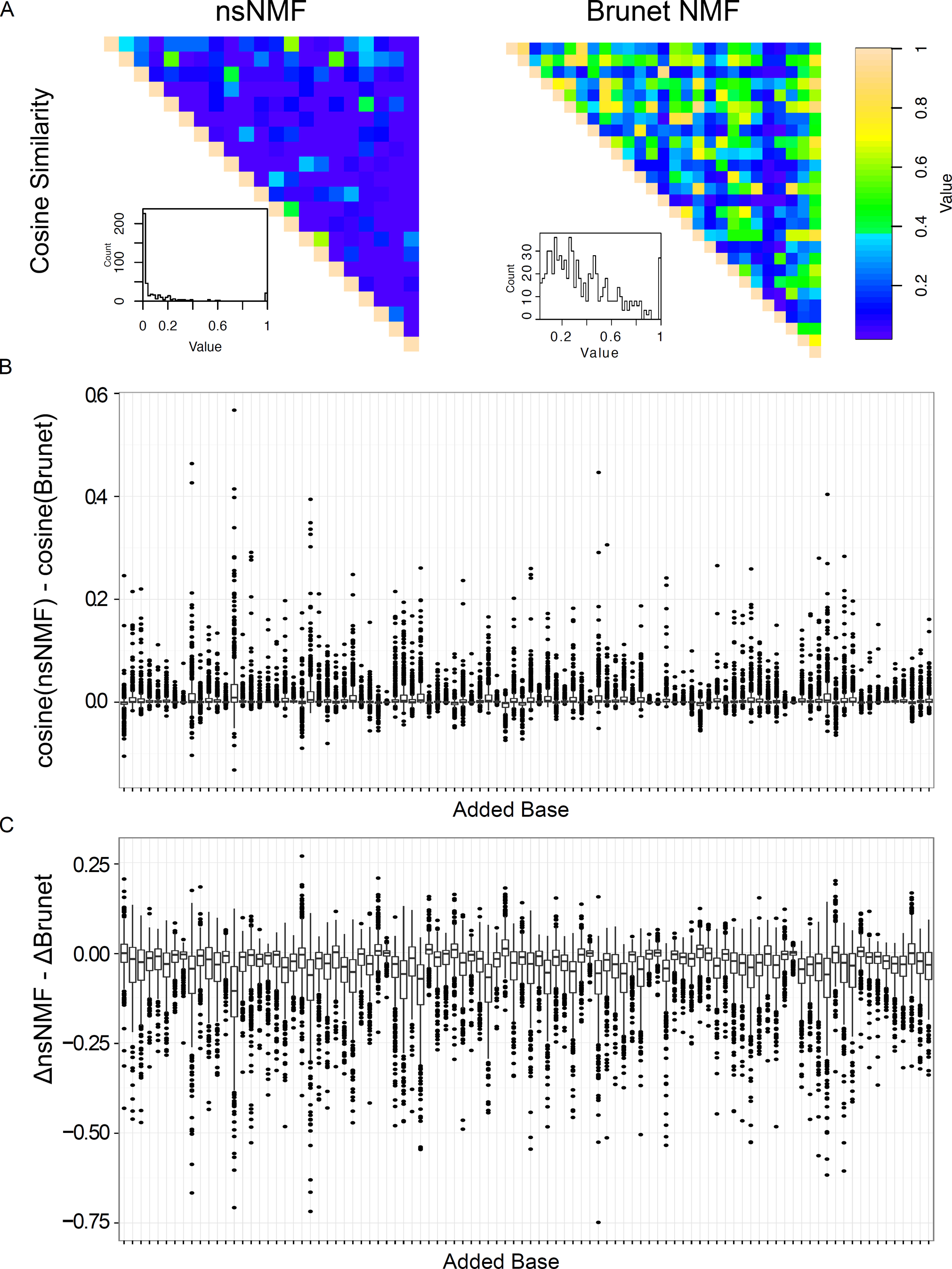
Similarity analysis of the 21 signature nsNMF solution compared with the Brunet solution. A) The cosine similarity for each signature pair in the respective studies is shown. Each panel is accompanied by a histogram of correlation values. Color is scaled from purple (0) to peach (1). B) Stability comparison of cosine similarity of mutation signature solutions simulated by adding a single variant to each sample in the Totoki liver cancer dataset and comparing the perturbed mutation set to the original. Values above zero indicate indicate more similarity from nsNMF compared to Brunet. C) Stability comparison of absolute deflection of perturbed verses original mutation signature solutions (see Supplemental Information). Values below zero indicate more stability from nsNMF compared to Brunet.

**Figure 2:**
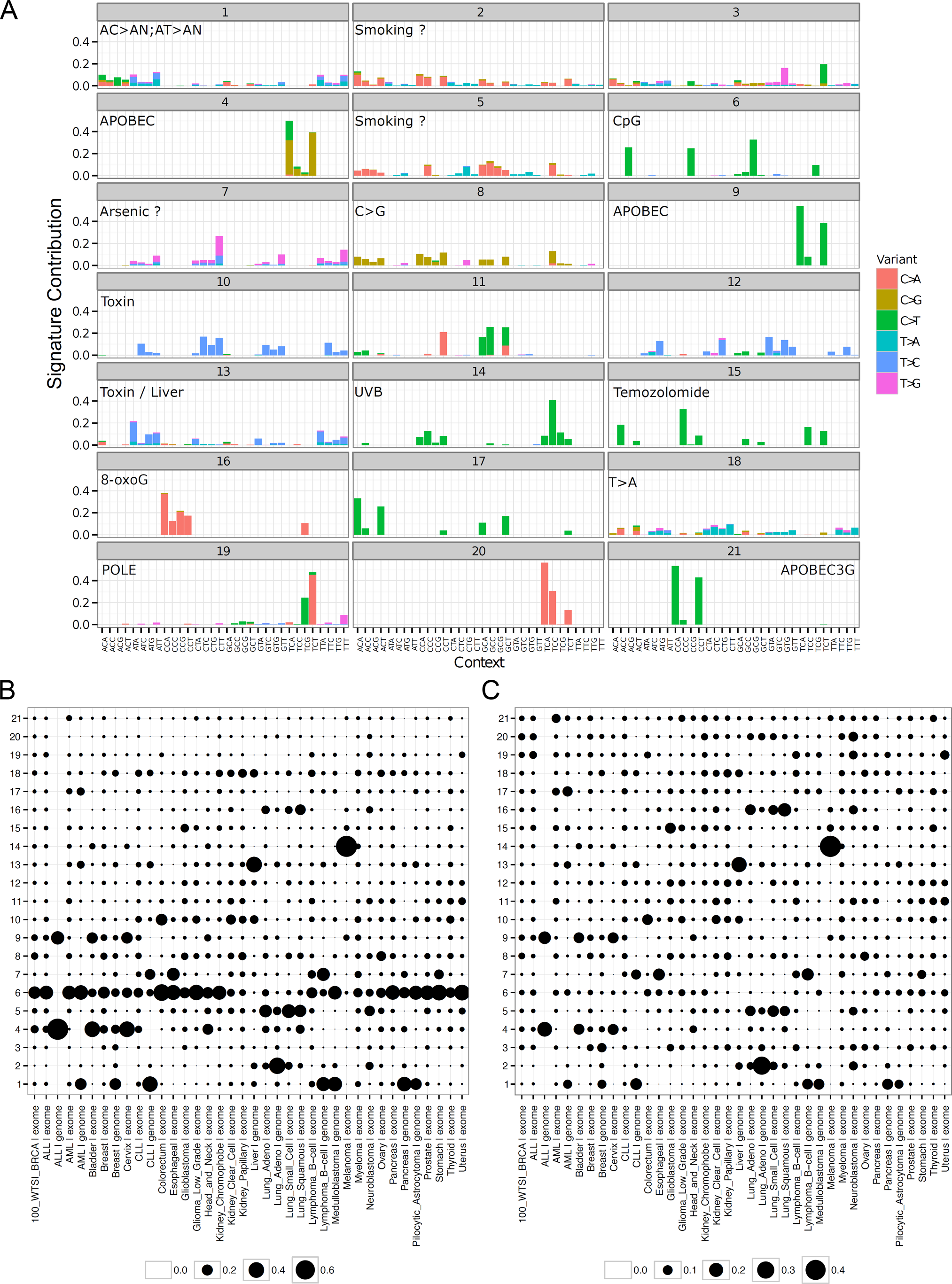
Mutation signature solutions in cancer. A)Mutation spectra for each of the 21 signatures is shown. Trinucleotide contexts are along the x-axis and relative contribution along the y-axis. Specific base changes are indicated by color. B) Scaled contribution of each mutation signature across cancer types. Signature contributions sum to 1 for each cancer (column). C) Scaled and weighted contribution of each cancer to the mutation spectra by scaling the data in B by signature (row).

**Figure 3:**
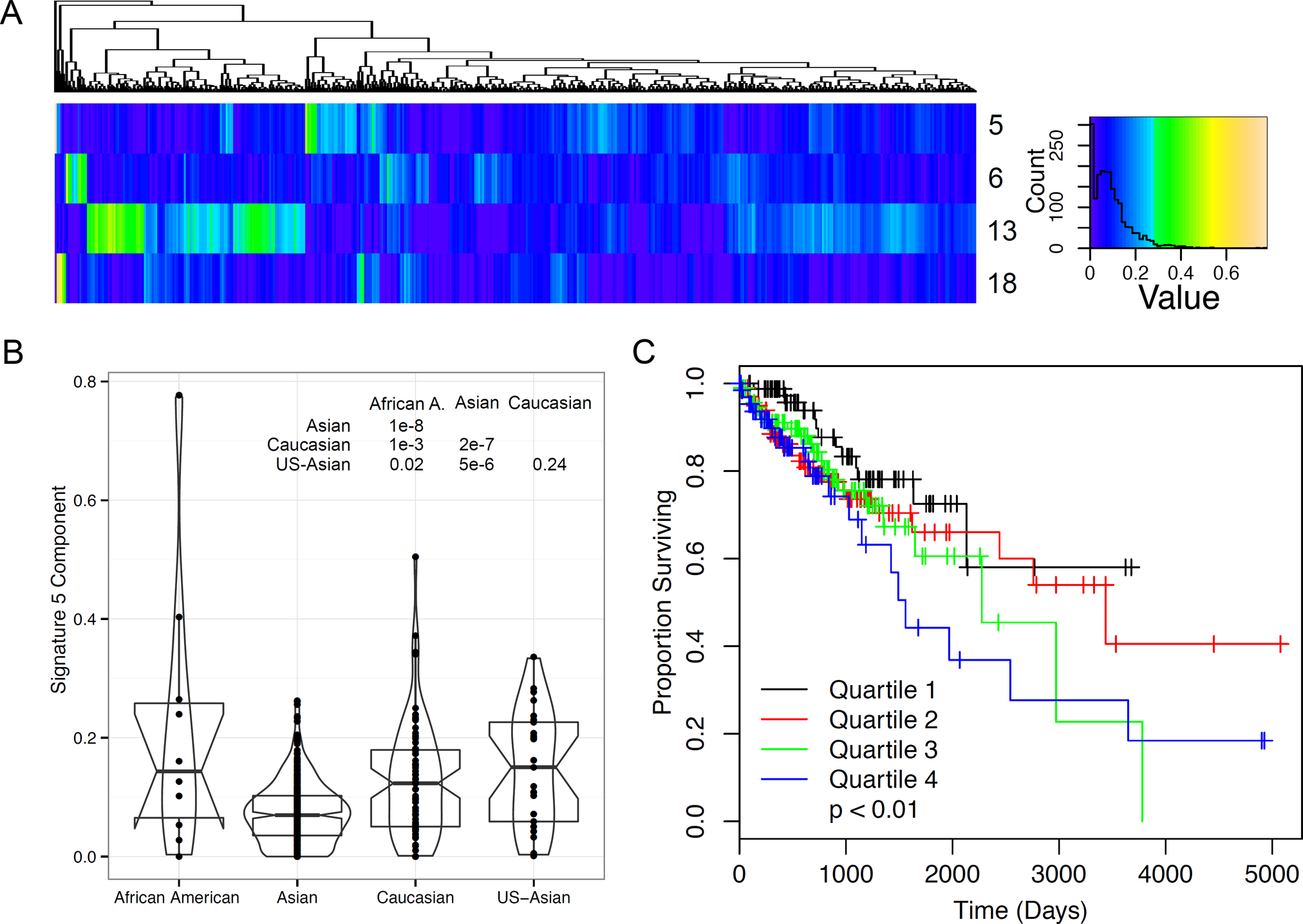
Mutation signature analysis from Totoki [14] dataset. A) Four signatures were found to be prevalent within the dataset, including signatures 5, 6, 13, and 18. A component histogram is shown to the right along with color scale bar. B) Mutation contribution of signature 5 by race, inset indicates the p-value of the indicated comparison, corrected for multiple comparisons using the Holm [20] method. C) Kaplain-Meier plot stratifying by quartiles of signature 5 proportion. Cox proportional hazards p<0.01.

## Discussion

nsNMF generated mutational spectra with superior mathematical properties compared to those previously reported. These spectra were associated with previously published biological mutation mechanisms. One should note that the nsNMF approach generates solutions with higher residual error compared to other methods. This effect is because of the reduction in noise within the solution set, which we consider an advantage for classification as demonstrated in stability analyses. These features reduce the impact of biological and sequencing noise on signature factorization yielding a better estimate of the actual mutational processes present, and thus, reducing noise in downstream analysis.

Our approach has also allowed us to extend previous work [5, 21, 6] beyond the source of the initial mutagen and yields insights into DNA damage repair. For example, we found that signature 1 was associated with whole genome sequence but was rarely present in exome data. We speculate that signature 1 represents a base substitution pattern that is efficiently repaired by transcription coupled repair but not by other mechanisms. The same may be the case for signature 2 in lung adeno carcinoma. We were also able to observe distinct trans-lesion synthesis pathways acting in the context of APOBEC signatures [8]. The capacity for nsNMF to identify very rare signatures is highlighted by the detection signature 21, an APOBEC3G mutation pattern [8, 9] which accounts for only 1.8% of the total mutation burden in the training dataset. The ability to detect these patterns is largely because of the special properties of nsNMF compared to standard NMF solutions. While NMF has been characterized as a “parts-list” there is no procedural mechanism for NMF to generate distinct signatures [4], therefore, NMF may not reveal minor signatures that nsNMF would detect.

**Figure 4:**
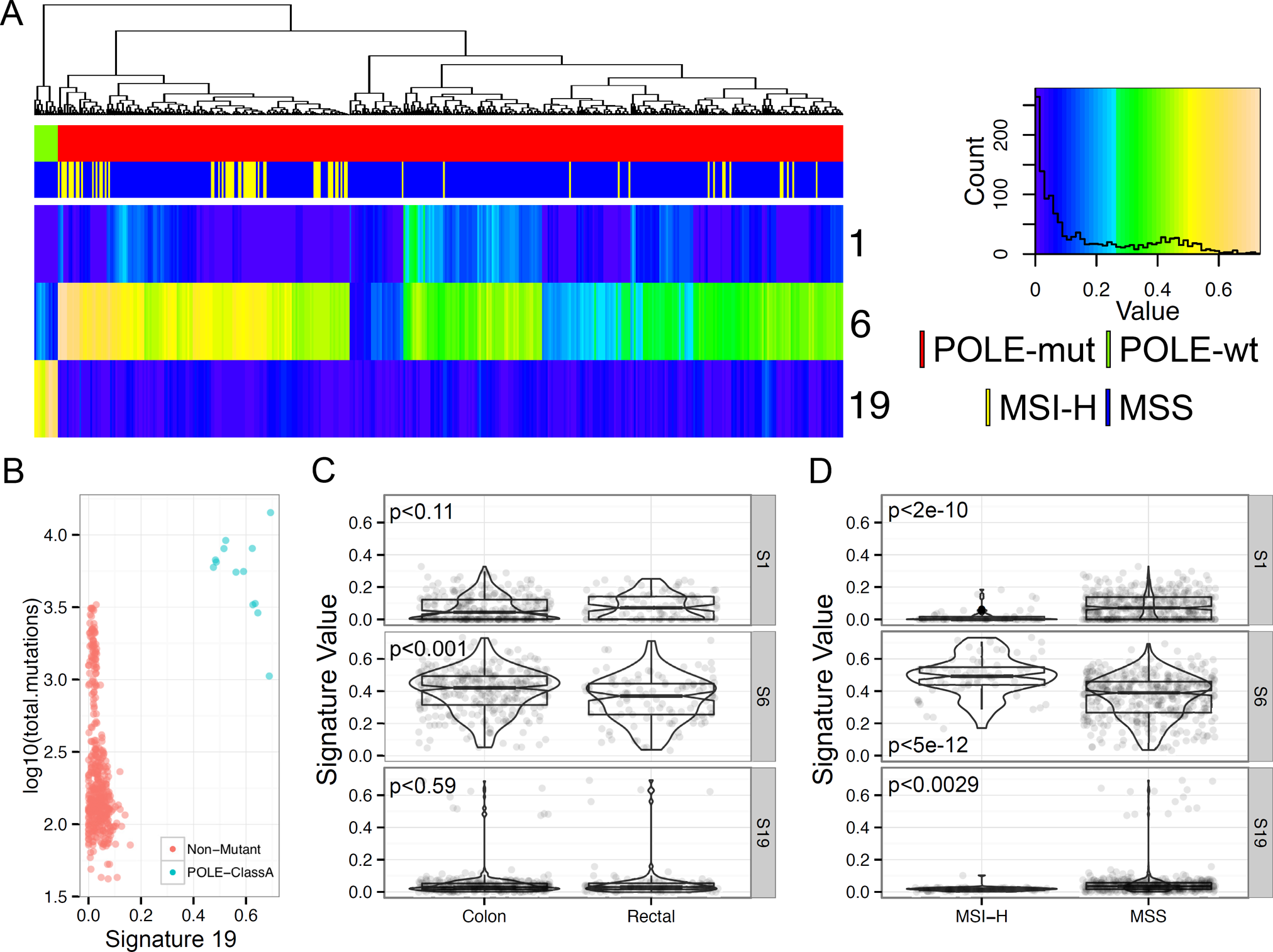
Mutation signatures in colorectal cancer. A) Clustering of signature coefficients in 449 colorectal cancer samples from the TCGA cohort. Three prevalent mutation signatures were noted; signatures 1, 6, and 19. The top track indicates POLE-exonulcease mutation status: POLE-mutant; red, POLE-wild type; green. The second track indicates microsatellite instability measures: MSI-high; yellow, MSS; blue. Count histogram of the coefficients for these signatures and color scale are shown to the right. B) Association of signature 19 with mutation counts and POLE-exonuclease mutation status. C and D) Comparison of signature coefficients for prevalent signatures by tissue type or MSI status. Insets represent p-values from a t-test.

We show two examples of the use of nsNMF-derived signatures for biological investigation. These nsNMF signatures make many types of additional investigation into the processes leading to cancer possible and represent a valuable resource to the genomics community. As such, we have made all of our classification resources freely available to the scientific community. We have ourselves used these signatures in several recent investigations including a study of cancers arising near the ampulla of Vader (ampullary cancer) where we identified signature 1 levels as a marker of poor outcomes (Gingras *et al.,* Cell Reports 2015, in press). We were also able to use signature 14 (UVB radiation exposure) to model the life history of Sezary syndrome cancers (Wang *et al.,* Nature Genetics 2015, in press). These studies highlight the utility of a thorough investigation of mutational exposures in the understanding of cancer.

## Methods

*Signature generation* Mutation data for the Alexandrov dataset were downloaded following the authors' instructions [5]. Duplicate samples were removed keeping the entries with the highest mutation counts. Mutation signatures for k = 15 to 25 were generated form the combined mutation count matrix using R and the NMF package [22] in the R statistical language [23]. All sessions for these fits are available from XXX (author note, XXX will be filled in at the time of publication when the data will be released). We performed 500 signature solutions with k = 21 signatures and the final model was selected based on minimal residual error in the model solution.

*Signature comparison* Signature comparison was performed by calculating the cosine similarity [24] for the pairwise combination of all solution vectors. Sparseness was calculated using algorithms in the NMF package [22].

*Signature stability* We tested the stability of the nsNMF by performing permutations using the Totoki dataset. We generated signature solutions for both the Brunet NMF and nsNMF signatures using unpermuted data. We then added to each sample a single mutation in a single variant context and then recalculated the signature solutions for this permuted dataset. We compared the cosine similarity between the original and permuted data and the absolute magnitude of difference between the original and permuted data. This process was repeated for each variant context.

*Signature analysis of Totoki dataset* Signature coefficients were generated for the Totoki [14] dataset using the reported mutations. Signature coefficients were scaled to sum to 1 for each subject and merged with clinical information. Survival analysis was performed using the survival package [25] in R.

*Signature analysis of TCGA CRC data* Mutation annotation files were generated for 449 COAD and READ samples and these mutations were used to generate signature coefficients. Signature coefficients were scaled to sum to 1 for each subject and merged with sample annotation including MSS/MSI status, POLE-mutation status and histological type.

*Other Analyses and Graphics* Figures were generated using utilities in the gplots [26] and ggplot2 [27] R packages. Data manipulation made extensive use of the reshape2 [28] and plyr [29] packages. All statistical analyses were performed in R [23].

All signature comparison analyses are reported in Supplemental information 1 and 2 as PDF and Sweave documents.

## Acknowledgments

The authors would like to thank Dr. Dmitry A. Gordenin for thoughtful discussion of cytosine deaminase DNA repair.

